# *Hapln1b* organizes the ECM to modulate *kit* signaling and control developmental hematopoiesis in zebrafish

**DOI:** 10.1101/2020.06.23.161406

**Authors:** Christopher B. Mahony, Corentin Pasche, Vincent Braunersreuther, Savvas N. Savvides, Ariane de Agostini, Julien Y. Bertrand

## Abstract

During early vertebrate development, hematopoietic stem cells (HSCs) are produced from a hemogenic endothelium located in the dorsal aorta, before they migrate to a transient niche where they expand, the fetal liver and the caudal hematopoietic tissue (CHT), in mammals and zebrafish, respectively. In zebrafish, previous studies have shown that the extracellular matrix (ECM) around the aorta needs to be degraded to allow HSCs to leave the aortic floor and reach blood circulation. However, the role of the ECM components in HSC specification has never been addressed. We show here that *hapln1b*, a key component of the ECM is specifically expressed in hematopoietic sites in the zebrafish embryo. Gain- and loss-of-function experiments all resulted in the absence of HSCs in the early embryo, showing that *hapln1b* is required, at the correct level, to specify HSCs in the hemogenic endothelium. Furthermore, we show that the expression of *hapln1b* is necessary to maintain the integrity of the ECM through its link domain. In addition, by combining functional analyses and computer modelling, we show that *kitlgb* interacts with the ECM, to specify HSCs. Overall, we have demonstrated that the ECM is an integral component of the microenvironment as it mediates specific cytokine signaling that is required for normal HSC specification.

## Introduction

The emergence of blood cells is a highly conserved process that consists of many different waves in distinct anatomical locations. The first emerging cells in zebrafish and mammals consist of primitive myeloid and erythroid cells ^1–4^. Definitive hematopoiesis is then initiated by the emergence of the transient erythro-myeloid precursors (EMPs), arising in the yolk sac in mice and humans ^1,2^, and in the posterior blood island in zebrafish embryos ^5^. Hematopoietic stem cells (HSCs) are then specified in the aorta-gonads-mesonephros (AGM) region where they form small intra-aortic clusters ^6–11^. HSCs are derived directly from the aortic hemogenic endothelium (HE), which is specified by the correct balance of several extrinsic factors, such as VEGF, Hedgehog, Notch, BMP and TGFβ signaling ^12–16^. Zebrafish HSC specification occurs between 32-60 hours post fertilization (hpf), from the HE in the dorsal aortal ^17,18^. Early HE specification in zebrafish is initiated by the expression of *gata2b* ^19^ followed by *runx1* ^12^. Zebrafish HSC specification is dependent upon inflammatory cytokines produced by neutrophils ^20–22^. Macrophage- and vascular-mediated extracellular matrix (ECM) degradation is also necessary to allow HSCs to migrate into the vein ^23,24^. HSCs will then migrate to the caudal hematopoietic tissue (CHT), where they interact with endothelial cells and significantly expand their initial number ^25,26^.

However, the exact mechanisms controlling HSC emergence from the HE and their expansion in the CHT remain to be fully characterized. We, and others, have previously published that these two processes are highly dependent upon cytokine signalling. In particular, in zebrafish, we showed the important and non-redundant roles of *oncostatin M* and *kit-ligand b(kitlgb)* in this process ^25,27^. Proteoglycans, a major component of the ECM, have been previously shown to interact with several hematopoietic cytokines (GM-CSF, IL-3 and Kitlg) and to maintain a close proximity between stromal cells, HSCs and cytokines in the niche ^28–30^. Here, we study the role of *hapln1b*, an ECM-associated protein, in contributing to this process. We focus on *hapln1b*, as it was previously shown to be expressed in the vasculature and in hematopoietic tissues ^31^. Furthermore, this gene was shown to be important for correct vascular development of the tail vasculature ^31^. In mammals there are *HAPLN1, 2, 3* and *4* genes ^32^, whereas in zebrafish there are *hapln1a, 1b, 2, 3* and *4* ^31^. *Hapln1* codes for a link protein, required to attach several chondroitin sulfate proteoglycans to the hyaluronic acid (HA) backbone (a ubiquitous glycosaminoglycan (GAG)) to make large, negatively charged, ECM structures ^33^. In mammals, *Hapln1* is a secreted ECM protein that stabilizes aggrecan-hyaluronan complexes and is required for correct craniofacial ^34^ and neocortex development ^35^. Loss of *HAPLN1* expression promotes melanoma metastasis but is also required to maintain immune cell motility ^36^. *HAPLN1* is also required to maintain lymphatic vessel integrity and reduce endothelial cell permeability preventing visceral metastasis ^37^. Knock-Out mouse studies have also revealed a role in maintaining perineural nets (PNNs), a hyaluronan backbone mesh-like network of proteins that surround neurons and regulate neuronal plasticity ^38^ as well as neural differentiation and development ^39–41^. Furthermore, PNNs are responsible for binding chemorepulsive molecules (such as *Semaphorin3a*) ^42^ as well as transcription factors, such as *Otx2* that is exchanged between different neural cells to enhance cortical plasticity ^43^.

Here we show that *hapln1b* is necessary for mediating *kitlgb-kitb* interactions, probably by regulating the ECM stiffness, to induce *runx1* expression in the HE during HSC specification in the zebrafish embryo. Gain- and loss-of-function of *hapln1b* resulted in defective hematopoiesis. Therefore, we conclude that this gene is required, at the correct dosage, for the specification of HSCs from the HE and their development after their emergence.

## Methods

### Zebrafish strains and husbandry

AB* (WT), along with transgenic and mutant strains were kept in a 14/10h light/dark cycle at 28°C ^44^. We used the following transgenic animals: *lmo2:eGFP_zf71_ ^45^, gata1:DsRed_sd2_* ^46^, *kdrl:eGFP_s843_* ^47^, *kdrl:Hsa.HRAS-mCherry_s896_* ^48^, *cmyb:GFP*_*zf169*_ ^49^, *globin:eGFP* ^50^, *sox10:mRFP*_vu234_ ^51^, *mpx*:GFP ^52^, *mpeg1:mcherry_gl23_* ^53^.

### Whole-mount In Situ hybridization (WISH) staining and analysis

WISH was performed on 4%PFA-fixed embryos at the developmental time points indicated. Digoxygenin labeled probes were synthesized using a RNA Labeling kit (SP6/T7; Roche). RNA probes were generated by linearization of TOPO-TA or ZeroBlunt vectors (Invitrogen) containing the PCR-amplified cDNA sequence. WISH was performed as previously described ^54^. Phenotypes were scored by comparing expression with siblings. All injections were repeated three independent times. Analysis was performed using R or GraphPad Prism software. Embryos were imaged in 100% glycerol using an Olympus MVX10 microscope. Oligonucleotide primers used for the production of ISH probes are listed in Table S2.

### Cell sorting and flow cytometry

Zebrafish transgenic embryos (fifteen to twenty per biological replicate) were incubated in 0.5mg/mL Liberase (Roche) solution and shaken for 90min at 33°C, then dissociated, filtered and resuspended in 0.9x PBS and 1% FCS. Dead cells were labeled and excluded by staining with 5nM SYTOX red (Life Technologies) or 300nM DRAQ7 (Biostatus). Cell sorting was performed using an Aria II (BD Biosciences) or a Biorad S3 (BioRad). Data was analyzed using FlowJo and GraphPad Prism.

### Quantitative real-time PCR (qPCR) and qPCR analysis

RNA was extracted using Qiagen RNeasy minikit (Qiagen) and reverse transcribed into cDNA using a Superscript III kit (Invitrogen) or qScript (Quanta Biosciences). qPCR was performed using KAPA SYBR FAST Universal qPCR Kit (KAPA BIOSYSTEMS) and run on a CFX connect Real time system (Bio Rad). All primers are listed in Table S1. Analysis was performed using Microsoft Excel or GraphPad Prism.

### Synthesis of full-length mRNA and microinjection

PCR primers to amplify cDNA of interest are listed in Table S2. *Kitlgb* and *kitlga* mRNA was synthesized and injected as previously described ^25,27^. mRNA was reverse transcribed using mMessage mMachine kit SP6 (Ambion) from a linearized pCS2+ vector containing PCR products. Following transcription, RNA was purified by phenol-chloroform extraction. *Hapln1b* mRNA was injected at 200pg, unless otherwise stated.

### Imaging

WISH were imaged using an Olympus MVX10 microscope in 100% glycerol. Fluorescent images were taken with an Olympus IX83 microscope and processed using cell sense dimension software. All images were processed using Adobe Photoshop. Time-lapse imaging was obtained using a Nikon inverted A1r spectral confocal microscope and processed using Fiji and analysed using GraphPad Prism.

### Morpholinos

All morpholinos oligonucleotides (MOs) were purchased from Gene Tools and listed in Table S3. MO efficiency was tested using PCR from total RNA extracted from a pool of 8-10 embryos at 24hpf. *Hapln1b* full-length primers were used to test MO efficiency and are listed in Table S2. *Hapln1b* morpholino was injected at 3ng, unless otherwise stated. All MOs used in this study are splice blocking MOs.

### Structural modelling and electrostatic properties of *kitlga* and *kitlgb*

Structural models for zebrafish kitlga and kitlgb were constructed based on a consensus approach employing homology-modeling, template-based structure modelling, and *ab initio* structure prediction as implemented in i-TASSER, Phyre2, and RaptorX ^55–57^. Vacuum electrostatic potential calculations were performed and displayed with the built-in module in the program PyMOL v. 2.3 (https://pymol.org/2/). Isoelectric point (pI) calculations were performed via the prot-pI server (https://www.protpi.ch).

### Glycosaminoglycan staining

Embryos were fixed in 4% formol (Biosystems) and embedded in paraffin (Haslab). 3μm sections were stained with 10mg/ml Alcian blue (Sigma) and counterstained with Mayer’s hemalun (Merck) to mark glycosaminoglycans.

## Results

### *hapln1b* is specifically expressed in early embryonic hematopoietic tissues

Previous studies have shown that *hapln1b* was expressed in the vasculature at 24-28hpf, and that its loss-of-function resulted in abnormal angiogenesis ^31^. However, its role during embryonic haematopoiesis has never been investigated. In zebrafish, *hapln1b* is the only *hapln* family member to display a vascular and hematopoietic expression pattern (data not shown), despite retaining high sequence identity and conserved peptides capable of forming disulphide bonds (Figure S1A, B). Synteny, phylogeny and sequence identity analyses revealed that *hapln1a* and *hapln1b* originate from a duplication of the *HAPLN1* gene in mammals (Figure S2A, B). Even across these species, peptides capable of forming disulphide bonds within the link domain have been conserved (Figure S2C). We therefore focused our study upon *hapln1b* and first examined its expression pattern. We first established that *hapln1b* is initially expressed from around 12hpf and would therefore not be derived from maternal RNA ^58^ (Figure S3A). *hapln1b* was expressed between 20hpf and 26hpf along the aorta, in the developing CHT and in the hypochord, as previously described ^31^. Further analyses at 26hpf revealed that *hapln1b* was also expressed in the aorta and vein region, ventral to the notochord (Figure 1Ai). Between 30 and 36hpf the expression begins to decrease, becoming more localized to the CHT, before being completely restricted to the marginal fold at 48hpf (Figure 1A). Expression was also scored in the cardiac precursors at 36hpf and in possible cranial cartilaginous structures (Figure 1Aii, Aiii). By 4 and 5dpf expression was restricted to cranial structures and was absent from hematopoietic tissue (Figure 1B).

**Figure 1.**
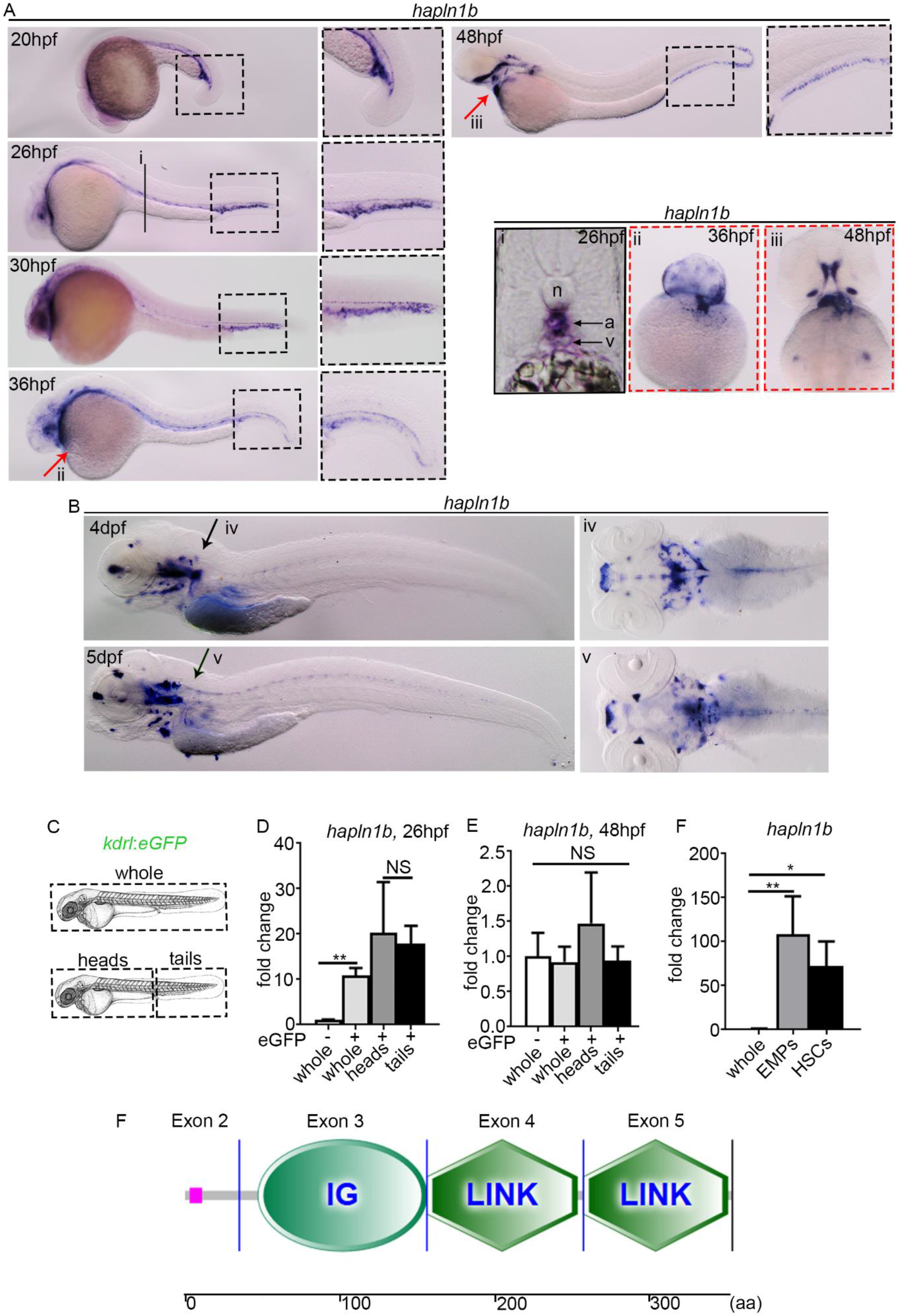
*hapln1b* is expressed by vascular cells and haematopoietic progenitors. (A) WMISH of *hapln1b* expression from 20hpf to 48hpf. (Ai) Section at 26hpf, displaying aorta (a) and vein (v) and notochord (n). (Aii and iii) ventral view at 36hpf and 48hpf. (B) WMISH of *hapln1b* expression at 4 and 5dpf. (Biv and v) Dorsal view at 4 and 5dpf. (C-E) Experimental outline and qPCR expression of *hapln1b* from FACS sorted endothelial cells. (F) qPCR expression of *hapln1b* in FACS sorted haematopoietic progenitors (EMPs and HSCs). All qPCR data is from biological triplicates. In D, two-tailed Student’s t-test, whole GFP− and GFP+, p=0.0035, heads and tails p=0.75. In E and F, analysis was performed by a one way ANOVA with multiple comparisons. In E, whole GFP− and GFP+, p=0.99, whole GFP− and heads, p=0.44, whole GFP− and tails, p=0.99. In F, whole and EMPs: p=0.0081, whole and HSCs: p=0.0449. (F) SMART (http://smartl.embl.de/) prediction of hapln1b protein structure. IG: immunoglobulin like-domain, Link: HA Link domain, aa: amino acids.

We then further analyzed the expression of *hapln1b* in different cell populations. We sorted endothelial cells from dissected *kdrl:eGFP* embryos at 26hpf and 48hpf (Figure 1C). As we found by wholemount in situ hybridisation (WISH), *hapln1b* was enriched in endothelial cells extracted from whole embryos at 26hpf (Figure 1D) but no enrichment was noted at 48hpf (Figure 1E). However, *hapln1b* was highly enriched in early EMPs (*lmo2:GFP;gata1:DsRed* double-positive cells at 28hpf ^5^) and nascent HSCs (*kdrl:mCherry*;*cmyb:GFP* double-positive cells at 36hpf ^17^), compared to whole embryos at 28hpf (Figure 1F), which showed a potential link between *hapln1b* and embryonic definitive haematopoiesis. Amino acid structural analysis (using SMART, http://smart.embl.de/) revealed that *hapln1b* contains an IG domain and two hyaluronic acid (HA) link domains (Figure 1F). We next investigated how this ECM protein could interfere with developmental hematopoiesis.

### *hapln1b* is required to specify HSCs from the Hemogenic Endothelium

To further investigate the role of this gene in hematopoiesis, we injected a splice blocking morpholino (MO) for *hapln1b* which efficiently reduced mRNA levels (Figure S3B, C). Inhibiting *hapln1b* expression did not affect *gata2b* expression, the earliest known marker of HE specification ^19^ (Figure 2A). However, *runx1* and *cmyb* expression was robustly decreased in the aortic region (Figure 2A). Accordingly, additional downstream expression of *cmyb* at 4dpf in the CHT and *rag1* at 4.5dpf in the thymus were also decreased (Figure 2A, B, and C). To further validate the specificity of our MO, we then rescued the loss of *runx1* in *hapln1b* morphants by co-injecting with *hapln1b* mRNA (Figure S3D). *hapln1b*-morphants displayed normal HSC specification, although in a number of cases the formation of the CHT was perturbed as seen in the “mild” cases, as previously described ^31^ (Figure 2D). In a small number of embryos, the vasculature was severely affected as represented in the “severe” phenotype (Figure 2D). We observed no change in the emergence of primitive macrophages or red blood cells as marked by *mfap4* and *gata1* at 24hpf (Figure 2E), respectively. We did however note a decrease in primitive circulating neutrophils, as marked by *mpx* (Figure 2E), although neutrophils were still present on the yolk sac. This change in distribution may be due to a loss of function of the vasculature that we observed in Figure 2D. Loss of *hapln1b* thus results into a specific loss of HSCs but does not impact primitive haematopoiesis. We then investigated the effect of *hapln1b* overexpression on embryonic haematopoiesis.

**Figure 2.**
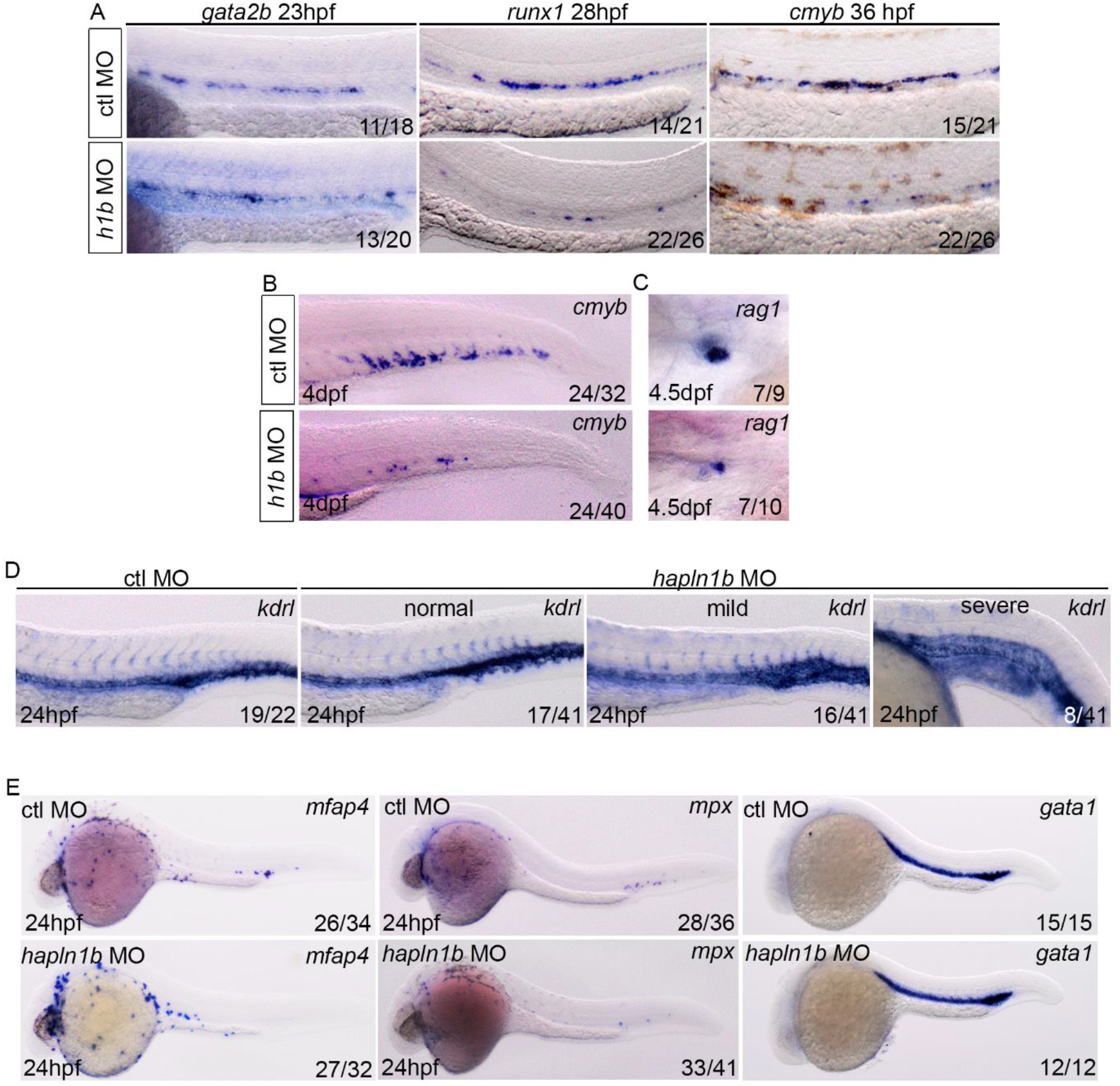
*hapln1b* is necessary for the specification of HSC from the hemogenic endothelium. (A) *gata2b, runx1* and *cmyb* ISH in control MO, or *hapln1b* MO injected embryos (injected at 3ng throughout, unless stated). (B) ISH expression of *cmyb*, in control MO, or *hapln1b* MO injected embryos. (C) *rag1* expression in the thymus after control MO or *hapln1b* MO injection. (D) ISH expression of *kdrl* in control MO, or *hapln1b* MO injected embryos. *Hapln1b* morphants displayed three phenotypes: normal, mild and sever. (E) ISH expression of *mfap4*, *mpx* and *gata1* in control MO, or *hapln1b* MO injected embryos. H1b: *hapln1b*.

### *hapln1b* overexpression is sufficient to reduce HSC emergence and downstream survival

We next overexpressed *hapln1b* by injecting the full-length mRNA at the one-cell stage and analysed the effect on developmental haematopoiesis. *hapln1b* overexpression did not change vessel development or early HSC specification as marked by *kdrl* at 24hpf and *runx1* at 28hpf, respectively (Figure 3A). However, we noted a decrease in *cmyb* signal at 36hpf in the AGM region (Figure 3A). *hapln1b* overexpression also decreased *cmyb* at 48hpf in the CHT, suggesting that newly formed HSCs did not colonize this tissue. However, in the AGM, the *cmyb* signal was also absent, as in non-injected controls, indicating that HSCs were not lodged and were not unable to migrate from the aorta (Figure 3B). Accordingly, *rag1* staining in the thymus at 4.5dpf and *cmyb* in the CHT at 4dpf were also decreased (Figure 3C). To further analyse this loss of *cmyb* signal, we then imaged double-positive *kdrl*:mCherry;*cmyb*:eGFP embryos to examine HSC emergence form the HE ^17^. *hapln1b* overexpression significantly reduced the number of double positive nascent HSCs from the AGM region at 36 and 48hpf (Figure 3D, D’). As expected, the number of HSCs present in the CHT niche at 3 and 4dpf was also significantly reduced (Figure 3E, E’). To further examine this loss of *cmyb* at 36hpf from the HE we used time-lapse confocal microscopy to image double-positive *kdrl*:mCherry; *cmyb*:eGFP embryos to examine the endothelial-to-hematopoietic transition in more detail in the aorta. We imaged from 34hpf to 42hpf to observe HSC budding from the HE. We observed clusters of cells in the AGM region of control embryos, preparing to bud and enter circulation (Movie S1). However, in *hapln1b*-overexpressing embryos, almost no cells seemed to initiate budding. Occasionally, some *cmyb:GFP*+ cells were detected, but none of them ever underwent endothelial-to-hematopoietic transition (Movie S2). We therefore conclude that *hapln1b* overexpression is sufficient to prevent the correct HSC budding from the HE. We next investigated how this could be occurring by examining how *hapln1b* could alter the ECM.

**Figure 3.**
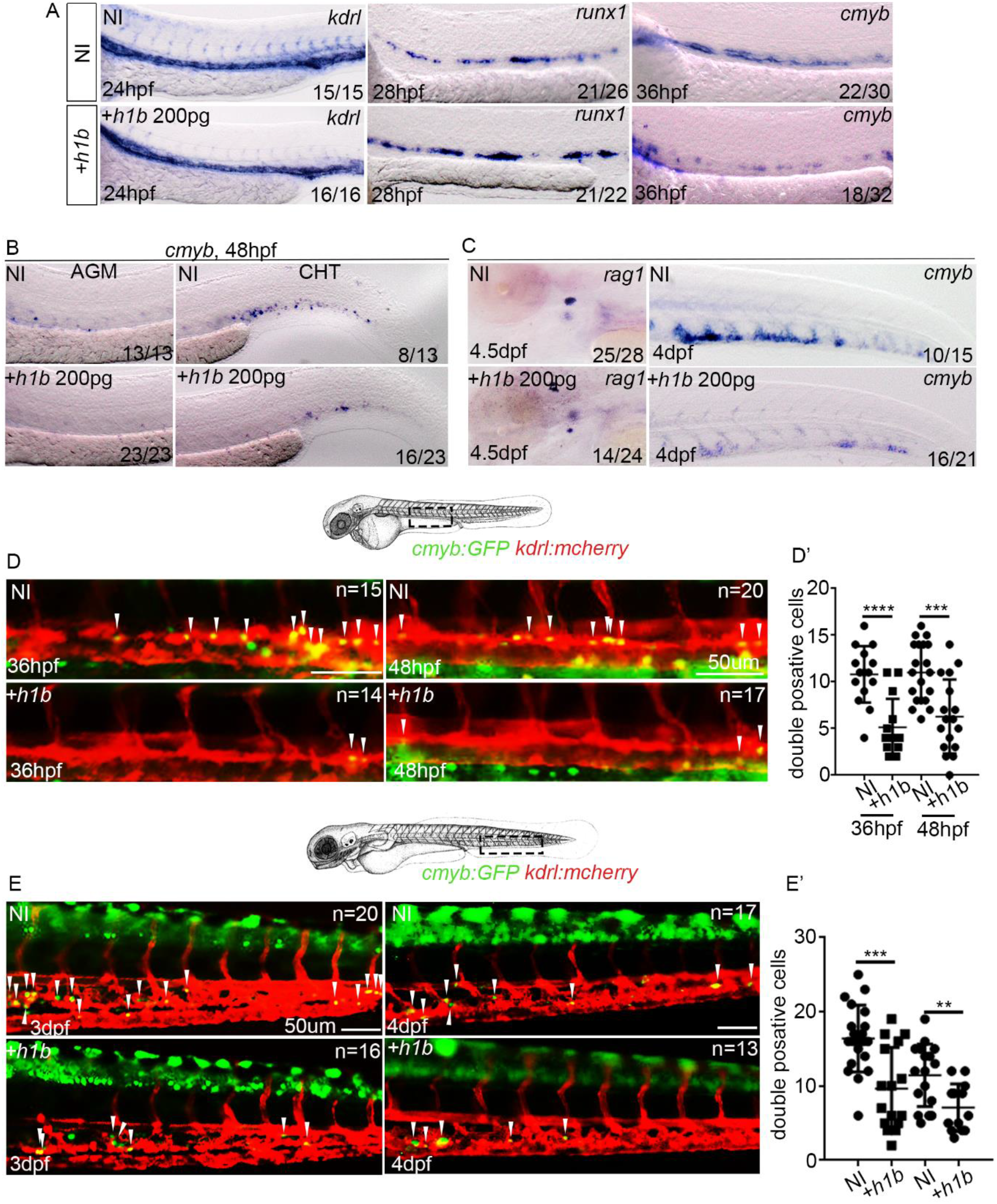
*hapln1b* overexpression is sufficient to inhibit HSC budding and development. (A) ISH expression of *kdrl*, *runx1* and *cmyb* in non-injected, or *hapln1b* mRNA injected embryos. (B, C) ISH expression of *cmyb* in non-injected, or *hapln1b* mRNA injected embryos. (D, D’) Imaging double transgenic *kdrl:mCherry/cmyb:GFP* embryos at 36 and 48hpf. Scale, 50μm. In D’, between NI and *+h1b* injected at 36hpf, p<0.0001 and at 48hpf p=0.0003. (E, E’) Imaging double transgenic *kdrl:mCherry/cmyb:GFP* embryos at 3dpf and 4dpf. Scale, 50μm. In E’, between NI and *+h1b* injected at 3dpf, p=0.0003 and at 4dpf p=0.004. NI: non-injected, +*h1b*: *hpln1b* mRNA injected.

### *hapln1b* modulates the density of the ECM around vasculature

The degradation of ECM by primitive macrophages is an important step for HSC emigration out of the aortic region ^23^. Therefore, we sought to confirm that *hapln1b* expression levels are capable of modulating the ECM directly. To achieve this, we examined glycosaminoglycans in the AGM and CHT region by staining embryo sections with alcian blue. Non-injected controls at 30hpf showed a high concentration of ECM surrounding the notochord, but little or no ECM around the artery and vein in the AGM (Figure 4A). *hapln1b*-moprhants showed less ECM staining, resulting in a change of the shape of the notochord, aorta and vein in the AGM, suggesting that the integrity of the ECM was altered (Figure 4A). In contrast, *hapln1b* mRNA overexpression resulted in increased ECM staining between the artery and vein and around vessels in the AGM (Figure 4A). Concerning the CHT, the artery and the venous plexus were clearly visible in controls morphants (Figure 4B). However, in *hapln1b-*morphants, the venous plexus was highly disrupted with no clear boundary between the artery and vein (Figure 4B). *hapln1b* mRNA overexpression resulted in a slightly distorted CHT with increased alcian blue staining and densely packed venous plexus (Figure 4B). All these results correlate with the importance of *hapln1b* in coordinating angiogenesis, as described previously ^31^ and could explain part of our results concerning developing haematopoiesis. We next examined the role that *hapln1b* plays in cytokine signalling to control this process.

**Figure 4.**
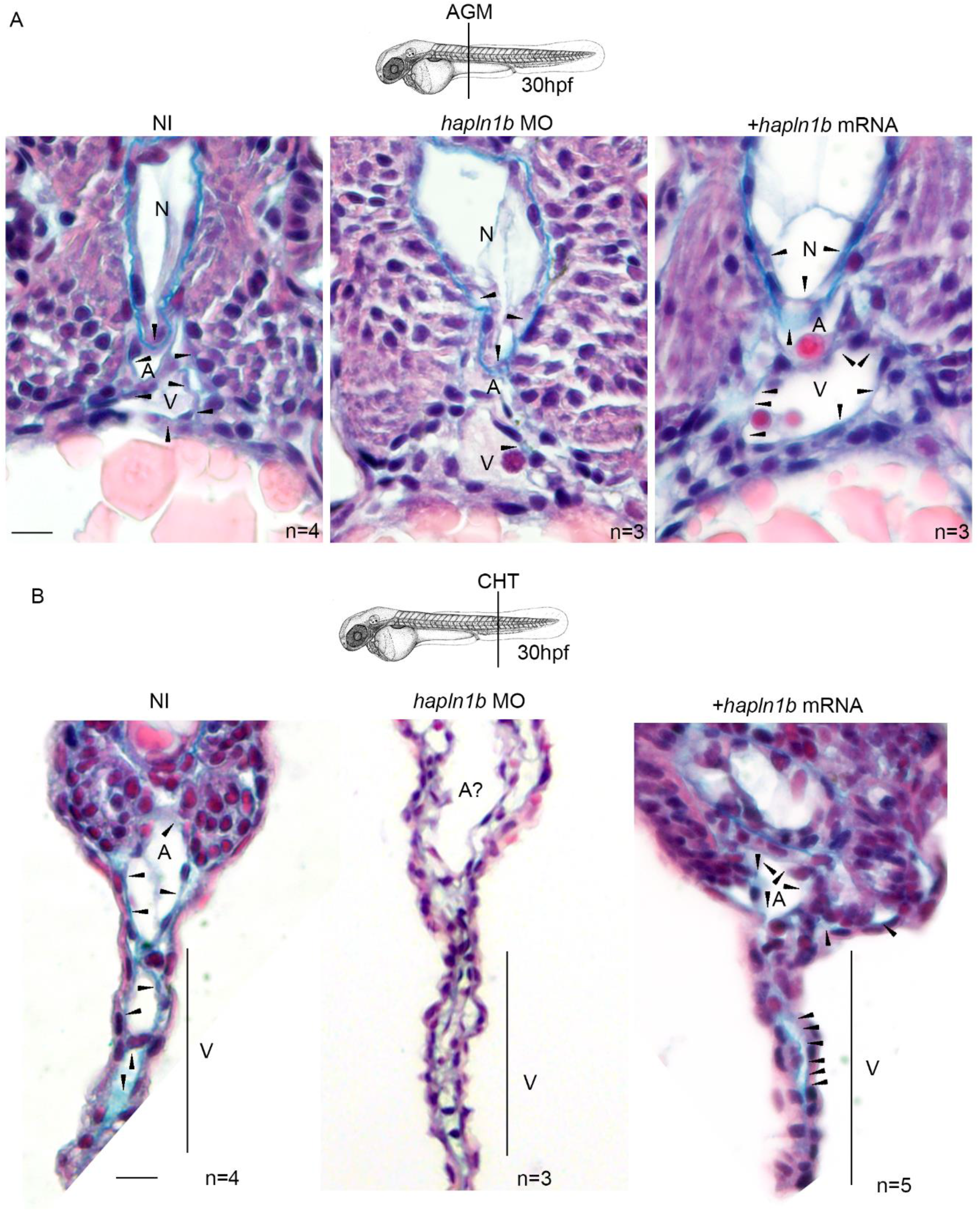
*hapln1b* is principally required for maintaining ECM integrity in the CHT. (A) Alcian blue and hemalun staining of glycosaminoglycans in the AGM region at 30hpf in NI. Black arrows indicate areas of ECM staining. (B) Alcian blue and hemalun staining of glycosaminoglycans in the CHT region at 30hpf. A= aorta, V=vein, N=Notochord. Scale bars, 10μm.

### *hapln1b* mediates *kitlgb-kitb* interactions that are required for proper HSC specification from the HE

We next attempted to decipher the mechanism by which *hapln1b* can affect haematopoiesis. Our previous studies indicated that HSCs express *kitb* and that *kiltgb* and *kitb* are required for *runx1* expression in the HE at 28hpf ^27^. As we found a similar decrease in *runx1* in *hapln1b*-morphants, we investigated a possible link between *kiltgb* signalling and *hapln1b*. Previous studies have indicated that *kitlgb* is expressed in the CHT region by ISH at 24hpf and is likely to exist more commonly as the membrane bound form, due to a loss of one of the two cleavage sites ^59^. Whereas *kitlga* would represent the more soluble form of this ligand as it retains the two cleavage sites ^59^. We also investigated the expression of *kiltgb* in many different FACS sorted cells and found no enrichment in ICM cells sorted from *globin*:GFP cells at 20hpf, primitive macrophages sorted from *mpeg1*:mcherry embryos at 24hpf, primitive neutrophils sorted from *mpx*:eGFP embryos at 24hpf and neural crest cells sorted from *sox10*:mRFP embryos at 26hpf (Figure S4 A-D). We also found no enrichment of *kitlgb* in endothelial cells sorted from dissected embryos at 19-20hpf (Figure S4E). However, we found a significant enrichment in tail endothelial cells at 26hpf (Figure S4F), corroborating with the previously published ISH expression pattern of *kitlgb* ^59^. We therefore concluded that *kiltgb* would be present in the AGM at low concentrations and would require an additional element to mediate effective interaction with its receptor to initiate *runx1* expression. We tested this hypothesis by attempting to rescue the loss of *runx1* observed in *hapln1b*-morphants and the loss of *cmyb* observed in *hapln1b*-overexpressing embryos by injecting *kitlgb* mRNA.

To verify the specificity of our *kitlgb* injection, we also injected *kitlga* mRNA, which does not play any role in HSC specification and development ^27^. Injection of either *kitlga* or *kitlgb* alone, did not alter expression of *runx1* at 28hpf, as anticipated (Figure 5A, A’). Injection of the *hapln1b*-MO gave a similar loss of *runx1* at 28hpf (Figure 5A, A’). We then co-injected the *hapln1b*-MO along with *kiltgb* mRNA, which in 16/34 embryos resulted in a rescue of the *runx1* signal (Figure 5A, A’). As expected, *kiltga* could not rescue the loss of *runx1* in *hapln1b*-morphants (Figure 5A, A’). We then attempted to rescue the decrease of *cmyb* signal in the *hapln1b*-overexpressing embryos by injecting *kitlgb* mRNA. Compared to non-injected controls, *hapln1b* mRNA reduced *cmyb* expression at 36hpf as observed earlier (Figure 5B, B’). Whereas *kitlgb* injection alone, did not alter the *cmyb* expression at 36hpf (Figure 5B, B’), co-injection of both *hapln1b* and *kitlgb* mRNAs increased *cmyb* expression to control levels (Figure 5B, B’). Altogether, this data suggests that the dose of hapln1b controls the stiffness of the ECM, therefore controlling the accessibility of *kitlgb* to the HE.

**Figure 5.**
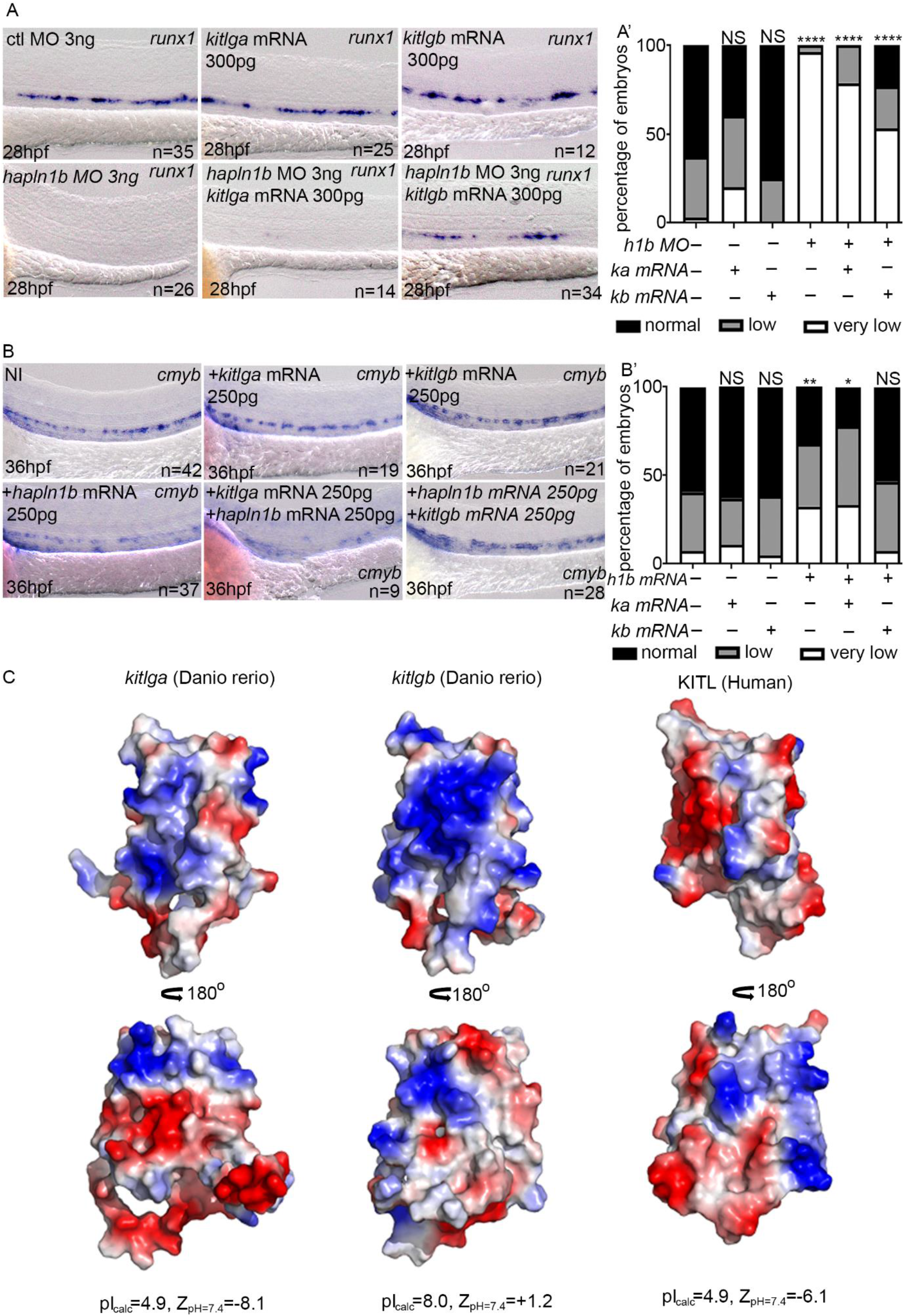
*hapln1b* interacts with *kitlgb* to maintain *kitlgb-kitb* interactions in the AGM. (A) ISH expression pattern of *runx1* in control morphants, *kitlga* mRNA injected embryos, *kitlgb* mRNA injected embryos, *hapln1b* morphants, embryos injected with both *hapln1b* MO and *kitlga* mRNA and embryos injected with both hapln1b MO and *kitlgb* mRNA. (A’) analysis of *runx1* expression. All analysis is compared to control MO. Data was analysed by fishers exact test using R. ctl MO vs +*kitlga*: p=0.05826, NI vs +*kitlgb*: p=0.7939, NI vs +*hapln1b*: p<0.0001, NI vs +*hapln1b*+*kitlga*: p<0.0001, NI vs +*hapln1b*+*kitgb*: p<0.0001. (B) ISH expression of *cmyb* in NI control embryos, *hapln1b* mRNA injected embryos, *kitlgb* mRNA injected embryos and embryos injected with both *hapln1b* and *kitlgb*. (B’) analysis of *cmyb* expression. All analysis is compared to NI control. Data was analysed by fishers exact test using R. NI vs +*kitlga*: p=0.8373, NI vs +*kitlgb*: p=1, NI vs +*hapln1b*: p=0.0051, NI vs +*hapln1b*+*kitlga*: p=0.036, NI vs +*hapln1b*+*kitgb*: p=0.8617. h1b, hapln1b. ka, kitlga. Kb, kitlgb. (C) Electrostatic potential of *KITL*, *kiltga* and *kitlgb* at biological pH.

As proteoglycans are negatively charged, we sought to examine the possible electrostatic compatibility of *kitlgb* with such a biological environment. In the absence of an experimentally determined structure for zebrafish *kitlgb* and *kitlga*, we leveraged available structural information at high resolution on human KITL (SCF) ^60,61^ and the related hematopoietic cytokines Flt3 ligand and colony stimulating factor 1 ^62–64^, to derive reliable models based on homology-based approaches and *ab initio* structure prediction.

Our analysis reveals that *Kitlgb* would be expected to display an overall positive electrostatic potential as manifested by a Z-potential value of +1.2 at physiological pH and as illustrated by extensive patches of positively charged amino acids at the protein surface (Figure 5C). Such physicochemical properties are in sharp contrast to zebrafish *Kitlga* and human *KITL*, which are pronouncedly acidic with very similar electrostatic properties characterized by strongly negative electrostatic Z-potentials at physiological pH. As proteoglycans are negatively charged, the overall positively charged *Kitlgb*, but not the negatively charged *Kitlga*, would mediate favourable interactions with the ECM in hemogenic and hematopoietic tissues, therefore activating the *Kitb* receptor, consistent with our findings.

### The link domain is necessary for *hapln1b* functions

Finally, we wanted to verify that *hapln1b* was interacting with HA by its link domain, to maintain ECM integrity. We cloned a truncated version of the *hapln1b* mRNA lacking exon 4 and 5 (*hapln1b*Δ4Δ5) that encode the HA link domain (Figure 6A). Compared to non-injected controls, *hapln1b*Δ4Δ5 mRNA injected did not reduce *cmyb* levels at 36hpf whereas wild-type *hapln1b* mRNA did (Figure 6B, B’), as shown earlier (Figure 3A). We therefore conclude that the link domain is necessary for *hapln1b* to exert its function, which is to assemble the ECM. We therefore propose the following model, where *kitlgb* is probably distributed along the aorta, by interacting with the ECM. The ECM structure is maintained by proteoglycans forming links with the HA backbone (mediated by the link domain of *hapln1b)*, allowing *kiltgb* to reach the AGM. A loss of *hapln1b* will prevent HA links to be formed and will prevent *kitlgb* distribution to the AGM, reducing *runx1* expression (Figure 6C). When *hapln1b* is overexpressed, HSC budding is reduced as the HA links are increased, resulting in a tight ECM that would also reduce *kitlgb* distribution to the AGM. Therefore, a lower concentration of *kitlgb* is present in the AGM would result in fewer HSCs budding and developing (Figure 6C).

**Figure 6.**
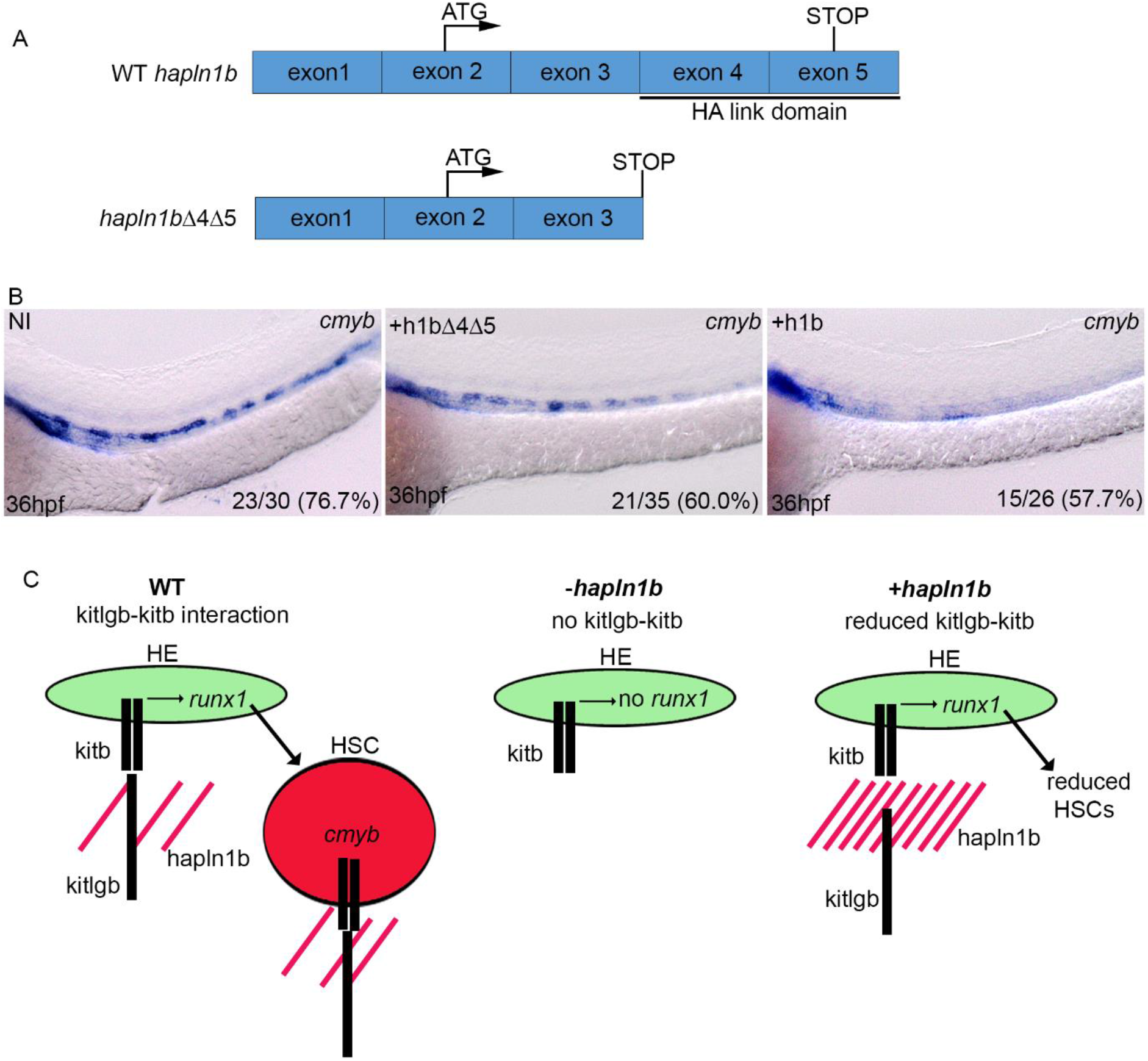
*hapln1b* maintains ECM integrity through the link domains. (A) Schematic representation of WT *hapln1b* and a truncated version that lacks exon 4 and 5 (*hapln1b*Δ4Δ5). (B) ISH expression of *cmyb* in NI control embryos, *hapln1b* mRNA injected embryos and *hapln1b*Δ4Δ5 injected embryos at 36hpf. (C) Summary of proposed model. In the WT situation, hapn1b helps *kitlgb* to interact with *kitb* in the AGM to permit runx1 expression and HSC formation. Loss of hapln1b results in a loss of kitlgb interacting with kitb, no *runx1* expression and no HSCs forming. Over expression of hapln1b results in a dense ECM and reduced kitlgb interacting with kitb, impairing HSC emergence and survival. HE: hemogenic endothelium.

## Discussion

We have shown that *hapln1b* is expressed along the embryonic endothelium prior to HSC emergence, before it becomes restricted to non-hematopoietic tissue at later stages (Figure 1). *hapln1b* is required, in the correct concentration, to specify HSCs from the HE and to maintain HSC budding and release into circulation. We found that loss of *hapln1b* results in a loss of *runx1* and loss of downstream HSC specification markers (Figure 2A-C). As proteoglycans can bind cytokines, it is possible that *hapln1b* is responsible for maintaining HA linking to proteoglycans to favor cytokine-receptor interaction. Our previous data has shown that *kitlgb-kitb* signalling is required for maintaining *runx1* expression and HSC specification in the HE ^27^. However, we, and others ^59^, have demonstrated that *kitlgb* is highly expressed in the posterior blood island but not the aortic HE region. When we modulate *hapln1b* expression, we are able to change the structure of the ECM as the loss of *hapln1b* affects the ECM/GAG deposition around axial vasculature, as well as in the CHT vessels and affects the overall CHT structure (Figure 4). The disruption of the CHT structure following *hapln1b* knockdown was also noted in a previous study ^31^. This, coupled with the fact that we are able to rescue *hapln1b* morphants with *kitlgb* mRNA injections (Figure 5A), lead us to conclude that *hapln1b* mediates HA linking with proteoglycans and is responsible for stabilising the ECM scaffold and distributing *kitlgb* to the HE (Figure 6C). Our data shows that the ECM likely plays a crucial role in maintaining the signalling environment in close proximity to the AGM to mediate HSC specification from the HE. Furthermore, we show an involvement of the ECM in the CHT, further suggesting that the ECM is a key player in maintaining the embryonic niche to help expanding HSCs. Although we have highlighted *kiltgb* as a key cytokine that interacts with the ECM, there are likely many others that contribute to the extrinsic signalling microenvironment. Fully characterizing and understanding these molecules will be an important step to improve the currently challenging derivation of HSCs from iPSCs.

In contrast to our loss-of-function assays, overexpression of *hapln1b* resulted in normal HSC specification as indicated by normal *runx1* expression (Figure 3A), but a reduction in HSC emergence and later expansion within hematopoietic tissue, as indicated by a reduction of *cmyb* expression (Figure 3B). As we found that HSC emergence was reduced, but never completely ablated, we reasoned that overexpression of *hapln1b* is likely to result in a denser ECM, reducing the accessibility of *kitlgb* to the hemogenic microenvironment. We confirmed this by rescuing HSC expansion in *hapln1b*-overexpressing embryos by overexpressing *kiltgb* mRNA (Figure 5B). Recent mouse studies have indicated that *Kitlg* is required throughout HSC development in the AGM region ^65^. This was further corroborated by additional studies showing that *c-kit* is expressed by proliferating HSCs in the AGM ^66^ and that *Kitlg* maintains HSCs in mouse AGM cultures ^67^. It is therefore likely that reducing the concentration of *kitlgb* in our *hapln1b*-overexpressing embryos restricts HSC emergence and decreases downstream survival and HSC maturation (Figure 6C). It is also possible that the ECM is too dense to allow HSC to bud and migrate to the CHT, as previous studies have indicated that ECM degradation is essential to HSC to enter blood circulation ^23,24^.

A similar role for *Hapln1* was also demonstrated in the PNNs in the mouse central nervous system, as it is required to maintain the HA mesh, surrounding neurons ^38^. These PNNs bind molecules to maintain neuronal plasticity and development, further implicating this gene in maintaining the ECM. Recent studies have identified that HSCs migrate to the fetal niche, the CHT, and expand their initial number in response to a number of cytokines ^24–26^. It is possible that many cytokine gradients and concentrations are maintained in the fetal niche by ECM components, such as *hapln1b*. Understanding these in more details will allow to fully characterizing HSC expansion providing alternative methods to improve HSC expansion *ex vivo*.

## Acknowledgements

J.Y.B. was endorsed by a Chair in Life Sciences funded by the Gabriella Giorgi-Cavaglieri Foundation and is also funded by the Swiss National Fund (310030_184814) and the “Fondation Privée des HUG”.

## Author contributions

SNS performed structural analysis of *KITL, kitlga* and *kitlgb*. VB and AA performed Alcian blue stainings on embryo sections. CBM and CP performed all other experiments. CBM and JYB designed the research and wrote the manuscript. SNS and AA edited the manuscript.

## Supplementary Tables

**Table S1.**
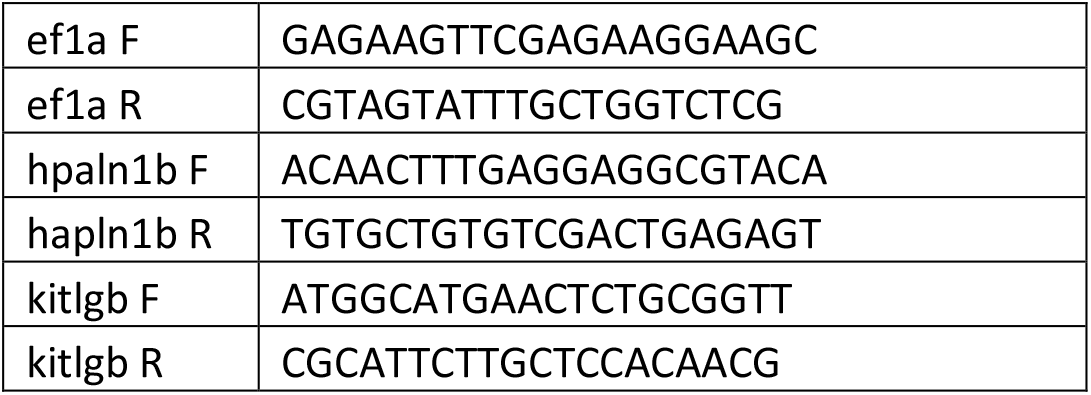
QPCR primers.

**Table S2.**
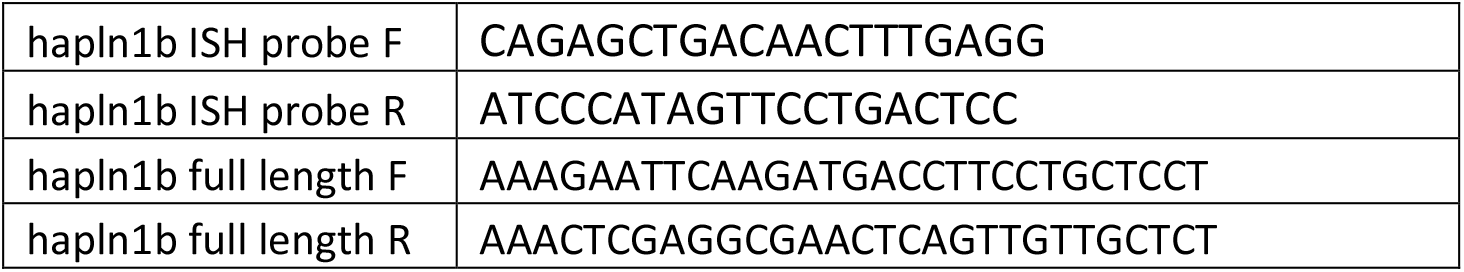
Cloning primers.

**Table S3.**
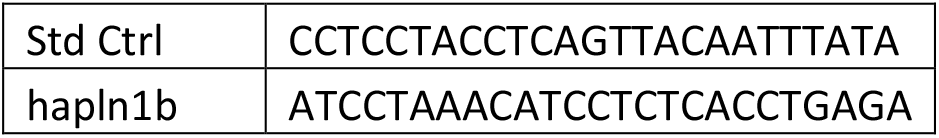
Mopholinos.

## Supplementary Figures

**Figure S1.**
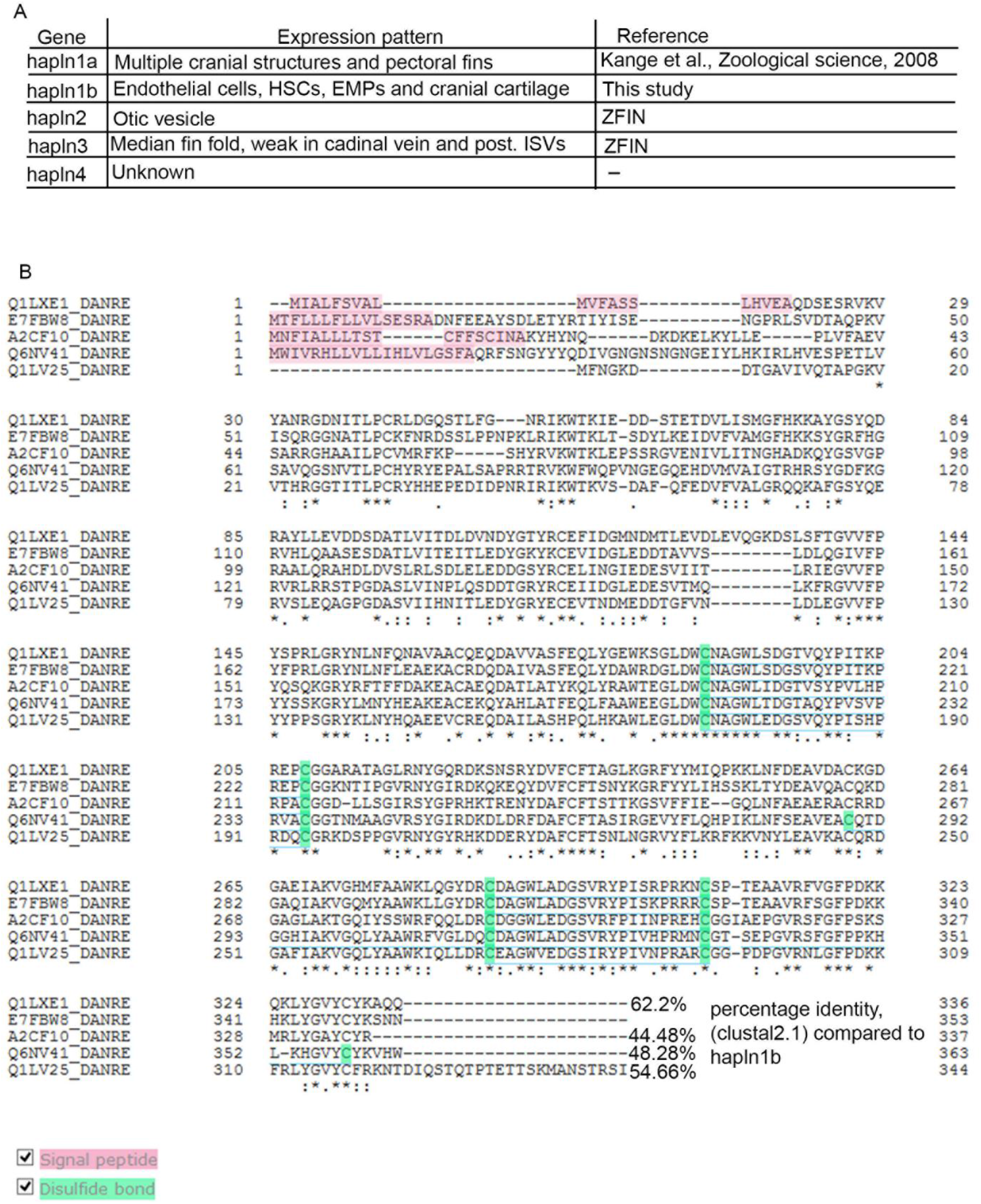
Comparison of hapln family member protein structures in zebrafish. (A) Summary of known expression pattern of other members of the *hapln* family in zebrafish. ZFIN: https://zfin.org/. (B) Amino acid sequence alignment (using uniprot) with highlighted signal peptide and disulphide bonds of zebrafish hapln family and percentage identity compared to hapln1b as calculated by clustal 2.1. Q1LXE1= *hapln1a*, E7FBW8= *hapln1b*, A2CF10=*hapln2*, Q6NV41=*hapln3*, Q1LV25=*hapln4*.

**Figure S2.**
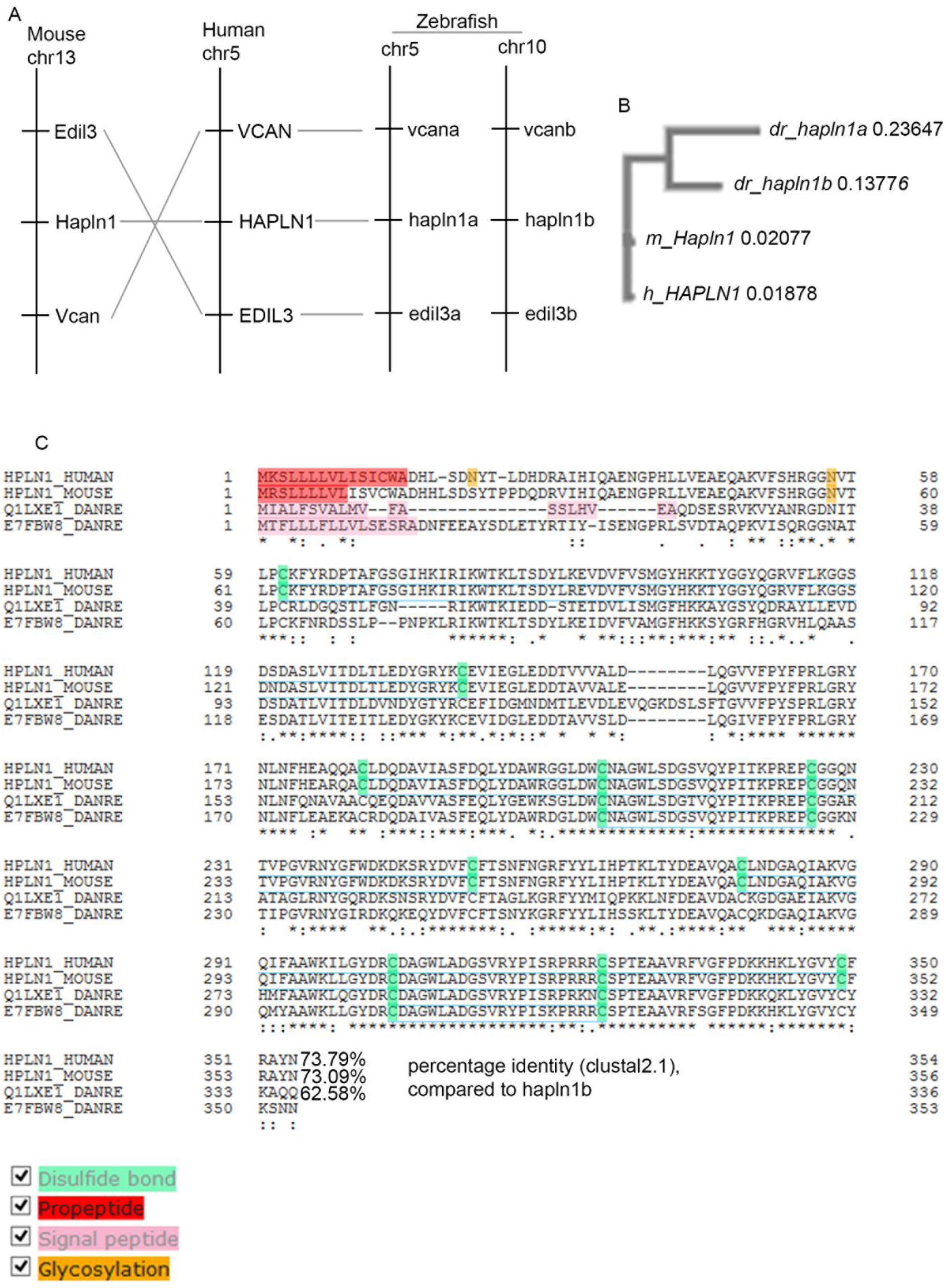
*hapln1b* and *hapln1a* arise from duplication of the mammalian orthologue. (A) Synteny analysis, showing duplication of the mammalian *hapln1* to give *hapln1a* and *hapln1b* in zebrafish. (B) phylogenetic analysis of *hapln1b* and its comparison to *hapln1a* in zebrafish (dr) and hapl1b in mice (m) and humans (h) as calculated by clustal 2.1. (C) Amino acid sequence alignment of zebrafish *hapl1b, and hapl1a, Hapl1 (mouse) and HAPLN1 (human)* with highlighted disulphide bond, propeptide, signal peptide and glycosylation site (analysed using uniprot). Percentage identity is compared to *hapln1b* as calculated by clustal 2.1. Q1LXE1= hapln1a, E7FBW8= hapln1b.

**Figure S3.**
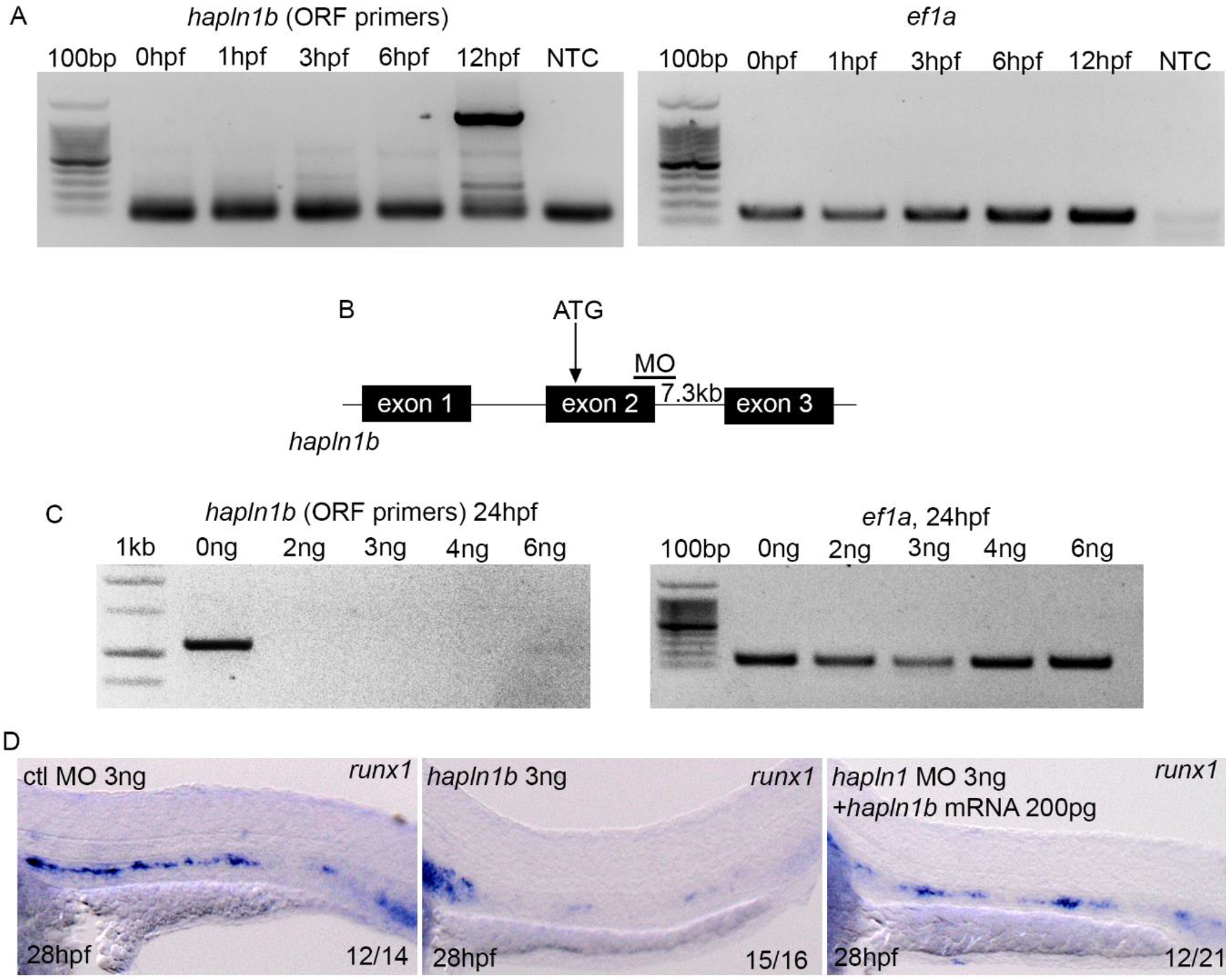
Validation of the *hapln1b* MO. (A) *hapln1b* expression by PCR from pools of 8-10 embryos extracted at the indicated timepoints. Primers for cloning full length hapln1b were used. *Ef1a* was used as a housekeeping control. NTC, no template control. (B) Schematic indicating MO target site, schematic is not to scale. (C) PCR after injection of the hapln1b MO at the indicated timepoints. RNA was extracted from pools of 8-10 embryos at 24hpf. *Hapln1b* full length primers used (ORF: open reading frame). *Ef1a* was used as a housekeeping control. (D) ISH expression of *runx1* in control morphants, *hapln1b* morphants and *hapln1b* morphants injected with *hapln1b* mRNA.

**Figure S4.**
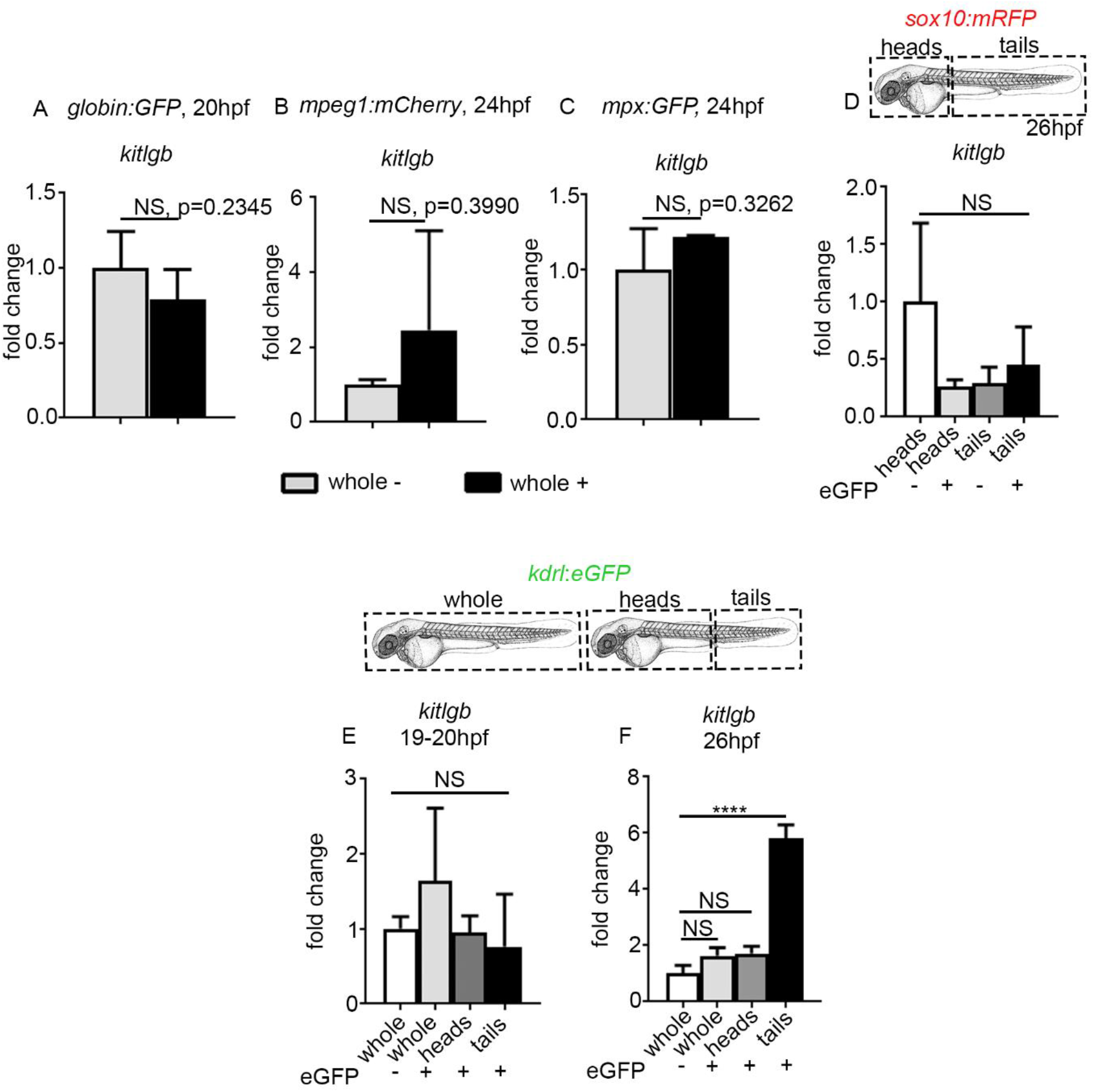
*kitlgb* is specifically expressed by caudal endothelial cells. (A-D) qPCR expression of *kitlgb* in either negative cells or positive cells from the indicated transgenic lines. In A-C, data is analysed using a two-tailed Student’s t-test, A: p=0.2345, B: p=0.3990, C: p=0.3262. In D, cells were sorted from dissection of heads or tails as indicated in the schematic, data was analysed using one-way anova. Heads− vs. heads+: p=0.1137, heads− vs. tails −: p=0.1294, heads− vs. tails+: p=0.2708. (E,F) Experimental outline and qPCR expression of *kitlgb* from FACS sorted endothelial cells from different dissections at either 19-20hpf or 26hpf. Data was analysed using one way anova. In E, whole− vs. whole+: p=0.4839, whole− vs. head +: p=0.9994, whole− vs. tail+: p=0.9338. In F, whole− vs. whole+: p=0.1300, whole− vs. head +: p=0.0866, whole− vs. tail+: p<0.0001.

